# *Pseudomonas aeruginosa* population dynamics in a vancomycin-induced murine model of gastrointestinal carriage

**DOI:** 10.1101/2024.08.19.608679

**Authors:** Marine Lebrun-Corbin, Bettina H. Cheung, Karthik Hullahalli, Katherine Dailey, Keith Bailey, Matthew K. Waldor, Richard G. Wunderink, Kelly E. R. Bachta, Alan R. Hauser

## Abstract

*Pseudomonas aeruginosa* is a common nosocomial pathogen and a major cause of morbidity and mortality in hospitalized patients. Multiple reports highlight that *P. aeruginosa* gastrointestinal colonization may precede systemic infections by this pathogen. Gaining a deeper insight into the dynamics of *P. aeruginosa* gastrointestinal carriage is an essential step in managing gastrointestinal colonization and could contribute to preventing bacterial transmission and progression to systemic infection. Here, we present a clinically relevant mouse model relying on parenteral vancomycin pretreatment and a single orogastric gavage of a controlled dose of *P. aeruginosa.* Robust carriage was observed with multiple clinical isolates, and carriage persisted for up to 60 days. Histological and microbiological examination of mice indicated that this model indeed represented carriage and not infection. We then used a barcoded *P. aeruginosa* library along with the sequence tag-based analysis of microbial populations (STAMPR) analytic pipeline to quantify bacterial population dynamics and bottlenecks during the establishment of the gastrointestinal carriage. Analysis indicated that most of the *P. aeruginosa* population was rapidly eliminated in the stomach, but the few bacteria that moved to the small intestine and the caecum expanded significantly. Hence, the stomach constitutes a significant barrier against gastrointestinal carriage of *P. aeruginosa,* which may have clinical implications for hospitalized patients.

**IMPORTANCE:** While *P. aeruginosa* is rarely part of the normal human microbiome, carriage of the bacterium is quite frequent in hospitalized patients and residents of long-term care facilities. *P. aeruginosa* carriage is a precursor to infection. Options for treating infections caused by difficult-to-treat *P. aeruginosa* strains are dwindling, underscoring the urgency to better understand and impede pre-infection stages, such as colonization. Here, we use vancomycin-treated mice to model antibiotic-treated patients who become colonized with *P. aeruginosa* in their gastrointestinal tracts. We identify the stomach as a major barrier to the establishment of gastrointestinal carriage. These findings suggest that efforts to prevent gastrointestinal colonization should focus not only on judicious use of antibiotics but also on investigation into how the stomach eliminates orally ingested *P. aeruginosa*.

## INTRODUCTION

In 2019, one in eight global deaths were attributable to bacterial infections (1). A handful of bacteria were responsible for half of these deaths, including *Pseudomonas aeruginosa,* which causes a wide range of healthcare-associated infections such as pneumonia, bloodstream infections, wound or surgical site infections, and urinary tract infections (1, 2). These infections are especially life-threatening for individuals who are hospitalized, immunocompromised, or have chronic lung diseases. In addition to being one of the leading causes of nosocomial infections (3), *P. aeruginosa* is also highly resistant to antimicrobial agents, making these infections difficult to treat (4). Some *P. aeruginosa* isolates are resistant to nearly all available antibiotics, including carbapenems, which has led to the CDC classifying multidrug-resistant *P. aeruginosa* as a serious threat (5).

*P. aeruginosa* is rarely part of the gastrointestinal (GI) microbiome of healthy individuals (4%) (6); however, it more efficiently colonizes patients in the intensive care unit (ICU) (10-55%) (6–8), with cancer (31%-74% of hospitalized patients) (9, 10), or in long-term care facilities (52%) (11). Importantly, GI carriage of *P. aeruginosa* is a key risk factor for subsequent development of infection (7, 12–14). As part of a prospective study, Gómez-Zorrilla *et al*. determined that after a 14-day stay in the ICU the probability of developing a *P. aeruginosa* infection was 26% for carriers versus 5% for noncarriers (13). In addition to the risk for infection, GI carriage can facilitate transmission of *P. aeruginosa* to other patients (15, 16). Thus, gaining a deeper understanding of *P. aeruginosa* GI carriage is crucial to prevent infections, manage rising rates of antibiotic resistance, and improve overall patient safety in healthcare settings.

While several animal models are available for the investigation of *P. aeruginosa* virulence and dissemination (17–24), fewer models have focused on GI carriage. In patients, antibiotic use has been correlated with an increased risk of *P. aeruginosa* GI colonization (12, 25–27). Previous studies have exploited this correlation to develop animal models that have provided important information on how *P. aeruginosa* establishes carriage. However, these models have relied on extended exposure to *P. aeruginosa* or the use of immunocompromised mice (18, 28, 29). It is estimated that around two-thirds of patients in the ICU are immunocompetent (30, 31). By using antibiotic pretreatment and immunocompetent animals, we aimed to develop an animal model of *P. aeruginosa* GI carriage that better mimics a typical ICU patient.

Here, we describe a murine model of *P. aeruginosa* GI carriage that is facilitated by the daily intraperitoneal (IP) injection of vancomycin for seven days and by a single dose of *P. aeruginosa* delivered via oral gavage. With this model, robust GI carriage was observed in both female and male mice, occurred with multiple clinical *P. aeruginosa* isolates, and persisted for up to 60 days. Additionally, to investigate the population dynamics of GI carriage, we used barcoded *P. aeruginosa* bacteria and determined that the stomach constituted a major barrier against GI carriage of *P. aeruginosa*. Bacteria that passed through the stomach were able to efficiently replicate in the small intestine and caecum, facilitating excretion of high numbers of *P. aeruginosa*. These barcoding experiments yielded interesting insights into the dynamics of *P. aeruginosa* GI carriage.

## RESULTS

### Vancomycin promotes gastrointestinal carriage of *P. aeruginosa*

Our goal was to develop a clinically relevant animal model that recapitulates the asymptomatic *P. aeruginosa* GI carriage observed in hospitalized patients. First, we tested the ability of the *P. aeruginosa* clinical isolate PABL048, delivered by orogastric gavage, to be carried in the GI tract of mice in the absence of antibiotic pretreatment. The extent of the carriage was assessed by fecal collection and plating on selective medium followed by CFU enumeration. Following pretreatment with IP phosphate-buffered saline (PBS), GI carriage of *P. aeruginosa* did not occur (Fig. 1A). Because antibiotic exposure correlates with an increased risk of GI colonization in patients (12, 25–27), we investigated the effect of IP vancomycin injection on the carriage of *P. aeruginosa* in mice. Vancomycin was chosen because it is one of the most commonly used antibiotics in the U.S. (32, 33) and does not have activity against *P. aeruginosa* (34). Three regimens of IP vancomycin treatment combined with orogastric gavage of *P. aeruginosa* were tested (all with a daily dose of 370 mg/kg of vancomycin – equivalent to a human dose of 30 mg/kg (35)). All three regimens supported the carriage of *P. aeruginosa* (Supplemental Fig.1). While vancomycin pretreatment for either 3 or 5 days prior to the orogastric gavage led to similar levels of GI carriage of *P. aeruginosa,* higher fecal burdens were observed when vancomycin injections were continued for two days after the orogastric gavage (Supplemental Fig.1). For all subsequent experiments, we therefore chose a regimen consisting of vancomycin on days −4 to −1, vancomycin and *P. aeruginosa* on day 0, and vancomycin on days +1 and +2 (Fig. 1B), which cumulatively corresponds to a typical 7-day course of vancomycin commonly prescribed to patients (36). When mice were treated with this regimen of vancomycin and challenged with 10^5.6^ CFU of the *P. aeruginosa* isolate PABL048, bacterial shedding averaged 10^6^-10^8^ CFU/g of feces during the first week (Fig. 1A). Although recovered CFU decreased somewhat during the second week post-inoculation, carriage levels remained between 10^3^ and 10^7^ CFU/g of feces. GI carriage of *P. aeruginosa* was similar in male and female mice (Fig. 1A).

**Figure 1:**
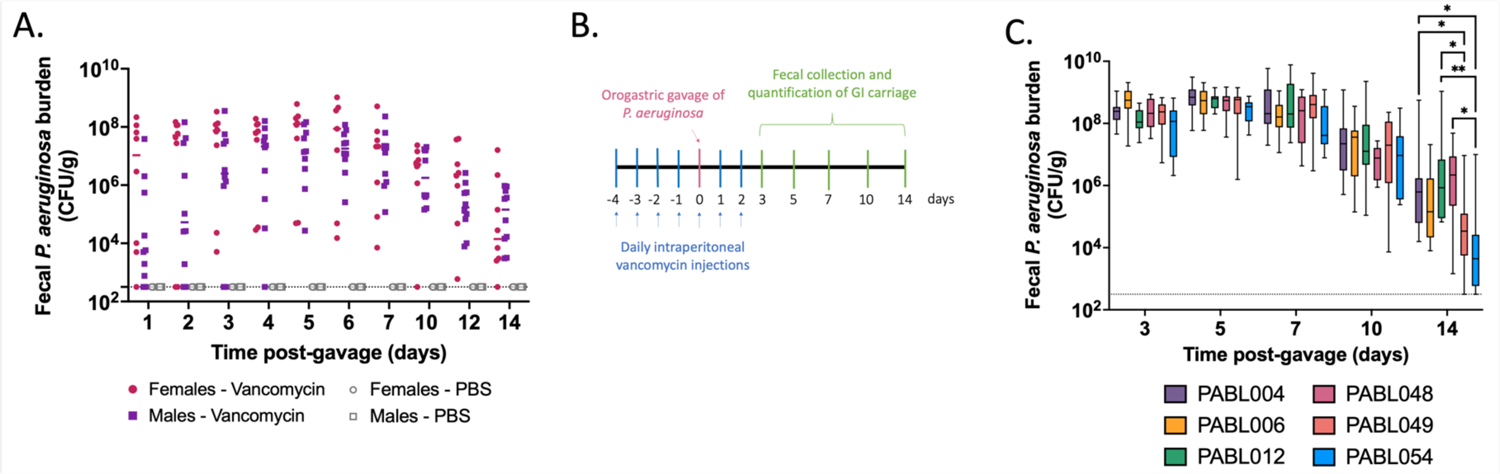
Murine model of *P. aeruginosa* GI carriage. (A) PABL048 fecal burden after an orogastric gavage with 10^5.6^ CFU. Male (square) or female (circle) mice received either PBS (gray symbols) or vancomycin (colored symbols) injections, and an orogastric gavage with 10^5.6^ CFU of PABL048. The experiment was performed twice for vancomycin treated mice (combined results are shown; n ≥ 8), and once for PBS treated mice (n = 5). Each symbol represents one mouse. Solid horizontal lines indicate medians. No significant differences in fecal CFU were found at any time point between male and female mice (multiple t-tests). (B) Timeline schematic of the model. Mice were intraperitoneally injected daily with vancomycin for 7 days at a dose of 370 mg/kg. On the fifth day of vancomycin treatment (day 0), mice received a defined dose of *P. aeruginosa* through orogastric gavage. On selected days, feces were collected to assess the extent of GI carriage, estimated by CFU counts. (C) Fecal burden of six clinical isolates of *P. aeruginosa* during GI carriage. Vancomycin and bacterial delivery (inoculum sizes: 10^5.4+/-0.2^ CFU PABL004, 10^5.6+/-0.3^ CFU PABL006, 10^6+/-0.2^ CFU PABL012, 10^5.4+/-0.1^ CFU PABL048, 10^6+/-0.1^ PABL049 or 10^6+/-0.1^ CFU PABL054) were performed as in B. Box plots are shown with boxes extending from the 25^th^ to 75^th^ percentiles, whiskers representing minimum and maximum values and lines indicating medians. Experiments were performed at least twice, and combined results are shown (n ≥ 10). *p ≤ 0.05, **p ≤ 0.01 (t-tests with Holm-Sidak correction for multiple comparisons). The dotted line indicates the limit of detection.

A substantial proportion of the *P. aeruginosa* genome is accessory (i.e., varies from strain to strain) (37). To examine whether these genomic differences allowed some strains of *P. aeruginosa* to establish higher or lower levels of GI carriage in this model, we individually inoculated mice with six clinical isolates: PABL004, PABL006, PABL012, PABL048, PABL049 and PABL054. These isolates are genetically diverse and exhibit differing levels of virulence in a bloodstream infection model (38) (Supplementary Table 1). Despite these differences, bacterial loads detected in the feces were similar for all strains over the first 10 days of the experiment (Fig. 1C). While more variability was observed on day 14, persistent carriage of all strains was detected. These results, taken together with the establishment of GI carriage in both sexes, demonstrate that vancomycin pretreatment produces a robust and reliable model to investigate *P. aeruginosa* carriage.

### GI carriage of *P. aeruginosa* does not cause GI inflammation

Carriage may be distinguished from infection by the absence of inflammation. To examine the vancomycin-treatment model represented true carriage, we performed histopathological analyses of the GI tract tissues. Mice received daily IP injections of vancomycin or PBS (day −4 to day +2) and were gavaged with either 10^7.1^ CFU of PABL048 or PBS (mock) on day 0. We chose to expose mice to a higher dose of bacteria than used in the previous experiments to maximize the possibility of observing inflammation. On day 3 post-orogastric gavage, mice were sacrificed and organs from the GI tract were harvested for histopathological examination. Organ sections were stained with hematoxylin-eosin (H&E) and screened for inflammation as evidenced by the presence of inflammatory cells or tissue damage. Neither of these were observed in any of the samples, and each section of the GI tract remained histologically normal (Fig. 2). Multifocal clusters of bacteria were observed adjacent to the mucosal surface of the stomach of all 3 mice that received IP PBS treatment and orogastric delivery of *P. aeruginosa.* These bacteria were primarily rod-shaped, compatible with *P. aeruginosa* morphology (39), but it was unclear whether they were dead or alive. Despite the presence of these bacteria in the stomach of PBS-treated mice three days after inoculation, *P. aeruginosa* bacteria were not cultured from their feces (Supplemental Fig. 2). Among mice treated with vancomycin prior to the bacterial inoculation, only one mouse exhibited bacteria adjacent to the surface mucosa of the stomach. While all mice that received vancomycin treatment and *P. aeruginosa* had feces that grew this bacterium (Supplemental Fig. 2), no bacteria were observed within the bowel walls of the small intestine, the caecum, or the colon of these same animals (Fig. 2), suggesting that *P. aeruginosa* remained in the lumen of the GI tract and did not invade the intestinal wall. In addition to these histopathological observations, mice exhibited no signs of systemic illness (e.g., decreased activity, ruffled fur) at any point during the experiment. Taken together, these results suggest that this is a model of GI carriage of *P. aeruginosa* rather than infection.

**Figure 2:**
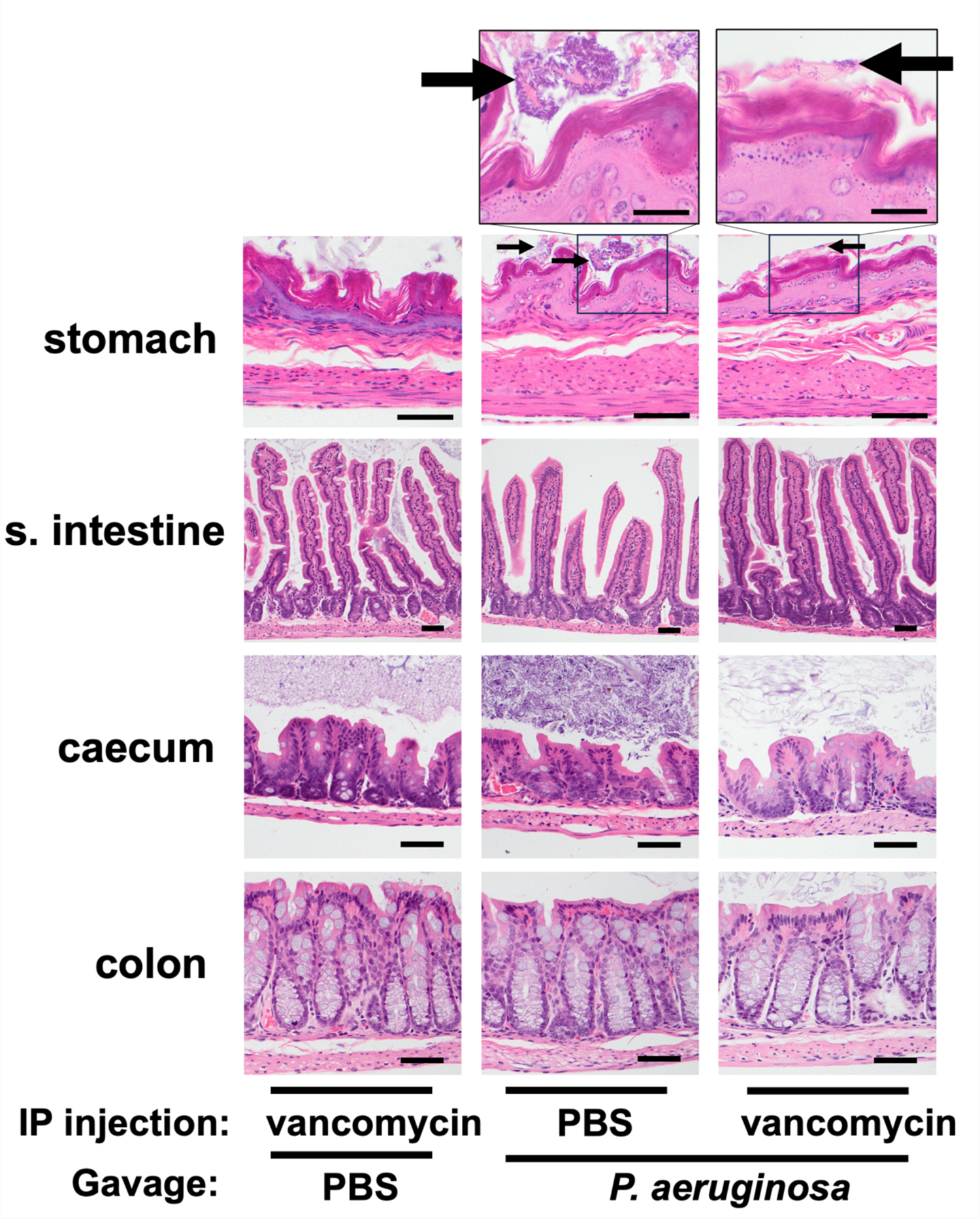
Tissue histology of the GI tracts of mice with carriage of *P. aeruginosa*. Hematoxylin-eosin staining of organ tissues collected at day 3 post-inoculation with either 10^7.1^ CFU of PABL048 or PBS (Mock). Prior to the orogastric gavage, mice received either vancomycin or PBS through intraperitoneal injections. Images in the bottom 4 rows were captured at a 400x magnification (bar = 100 µm). Images on the top row were captured at a 1,000x magnification (bar = 200 µm). (n = 3-4 mice/group). Arrows indicate intraluminal clumps of bacterial bacilli.

### *P. aeruginosa* bacteria remain largely within the GI tract

Since the intestinal tract has previously been identified as the main source of *P. aeruginosa* for the development of infections in immunocompromised patients (12, 40, 41), we assessed whether this model resulted in dissemination of *P. aeruginosa* to other tissues. Mice were orogastrically inoculated with 10^7.4^ CFU of PABL048, and *P. aeruginosa* CFU were enumerated from various organs at days 3, 7 and 14 post gavage. Most of the bacteria were detected in the organs of the GI tract, including the stomach, small intestine, caecum, colon, and feces (Fig. 3). Nevertheless, *P. aeruginosa* was occasionally detected in the gallbladder, spleen, liver, or lungs within the first 7 days post gavage. This suggests that, in this model, escape of bacteria from the gut, while very infrequent, did occasionally occur in the absence of observable signs of systemic illness. However, by two weeks post-inoculation, we did not observe bacteria in any systemic site of the mice. In summary, dissemination of *P. aeruginosa* from the GI tract is rare in this model of GI carriage.

**Figure 3:**
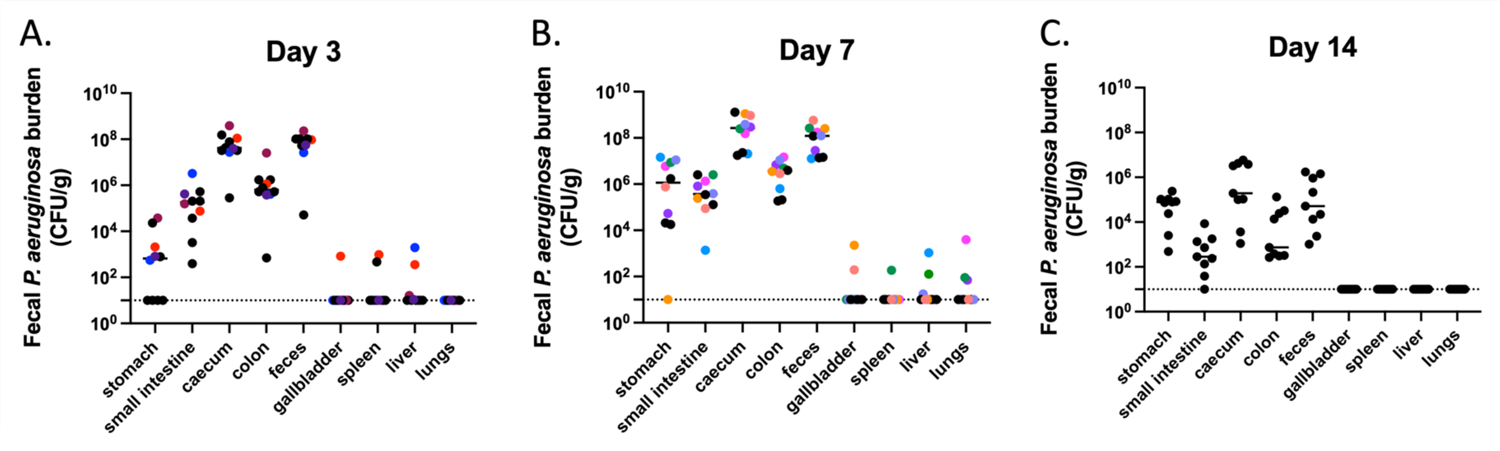
Dissemination of *P. aeruginosa* from the GI tract. *P. aeruginosa* burden in organs of mice carrying PABL048. Mice were sacrificed at (A) day 3 (n = 10), (B) day 7 (n = 10) or (C) day 14 (n = 9) post-orogastric gavage with 10^7.4+/-0.2^ CFU of PABL048, and bacterial CFU in the organs were enumerated by plating. The experiment was performed twice; combined results are shown. Each symbol represents one mouse. Solid horizontal lines indicate medians. The vertical dashed lines separate GI tract organs (left) from other organs (right). Symbols representing mice with dissemination from the GI tract are colored (one color/mouse). The horizontal dotted line indicates the limit of detection.

### *P. aeruginosa* establishes long-term carriage

To further characterize this model, we interrogated the duration of carriage following a single orogastric gavage of *P. aeruginosa.* When inoculated with 10^5.7^ CFU of PABL048, all mice carried *P. aeruginosa* in their GI tract for at least 10 days (Fig. 4). At day 60, 70% of all mice (7/10 females, 7/10 males) were still shedding *P. aeruginosa* from their GI tract. These results show that, in this model, *P. aeruginosa* establishes long-term GI carriage following a single exposure.

**Figure 4:**
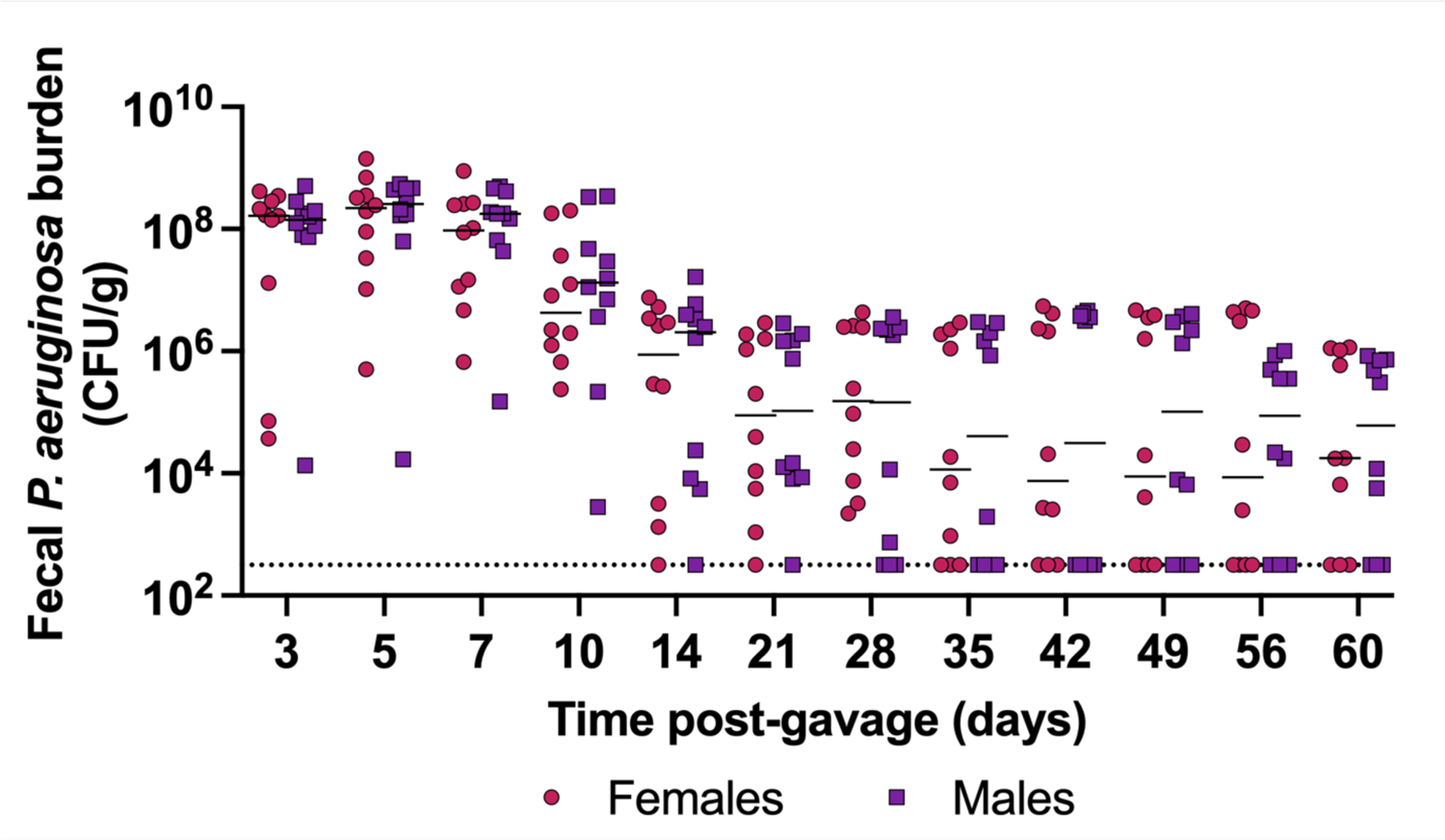
Long-term carriage of *P. aeruginosa* in the GI tract. PABL048 fecal burdens after an orogastric gavage with 10^5.7+/-0.3^ CFU in male (purple squares) and female (red circles) mice. The experiment was performed twice; combined results are shown (n = 10). Each symbol represents one mouse. Solid horizontal lines indicate medians. No significant differences were found between male and female mice at any time point (multiple t-tests). The dotted line indicates the limit of detection.

### The stomach constitutes the main bottleneck of GI carriage

Investigation across the segments of the GI tract indicated that not all tissues supported the same levels of *P. aeruginosa* carriage (Fig. 3). Thus, we sought to examine the GI carriage dynamics following orogastric inoculation with *P. aeruginosa.* In particular, we wanted to identify which segments of the GI tract contributed to population bottlenecks or supported bacterial expansion in this model. The sequence tag-based analysis of microbial populations (STAMP) technique (42), which relies on the generation of a bacterial library with insertions of short, random nucleotide DNA tags into a neutral site of the chromosome, is ideal for this purpose. Animals are inoculated with this library, and barcode frequency and diversity at different locations and times post inoculation are interpreted using the refined framework of STAMP (known as “STAMPR”) (43). This analysis estimates the size of the founding population (N_s_), defined as the number of bacterial cells from the inoculum that successfully passed through physical, chemical and immune barriers in the host to establish the population at the site of infection. A low N_s_ value (a small number of unique barcodes) is indicative of a tight bottleneck, while a high N_s_ value (a large number of unique barcodes) is reflective of a wide bottleneck. Comparison of CFU obtained from a tissue with the N_s_ value provides insight into the extent of bacterial expansion; for example, high CFU could be obtained from 1) a wide bottleneck followed by little bacterial replication or 2) a tight bottleneck followed by extensive replication of a small number of founders.

We applied STAMP to the vancomycin-treated mouse model. We used a previously generated barcoded library in the *P. aeruginosa* clinical isolate PABL012 (∼6,000 unique tags, each ∼30 bp) that had been validated by Bachta and colleagues (designated “PABL012_pool_”) (44). Note that fecal shedding of PABL012 was similar to that of PABL048 in the vancomycin-treated mouse model (Fig. 1C). Because we observed stable GI carriage of PABL012 between days 3 and 7, we deduced that major steps of GI carriage establishment were likely to occur within the first 3 days following inoculation. Using the vancomycin-treated mouse model, we delivered 10^6.1^ CFU of the PABL012_pool_ library through orogastric gavage and collected and analyzed segments of the GI tract (stomach, small intestine, caecum, colon, and feces) at 24, 48, or 72 hours post-inoculation (hpi).

As previously observed with PABL048 (Fig. 3), all organs of the GI tract supported the carriage of PABL012_pool_ (Fig. 5A-D). The stomach was the organ with the largest variation in total CFU recovered (Fig. 5 A-D); while *P. aeruginosa* was no longer recovered from the stomach of some mice, others carried 10^4-5^ CFU. The caecum and the feces had the highest bacterial burdens during the first 3 days of GI carriage. Median CFU loads recovered from all sites were stable over the first 3 days (Fig. 5D), suggesting that GI carriage is established during the first 24 hours following *P. aeruginosa* delivery and maintained for the next two days.

**Figure 5:**
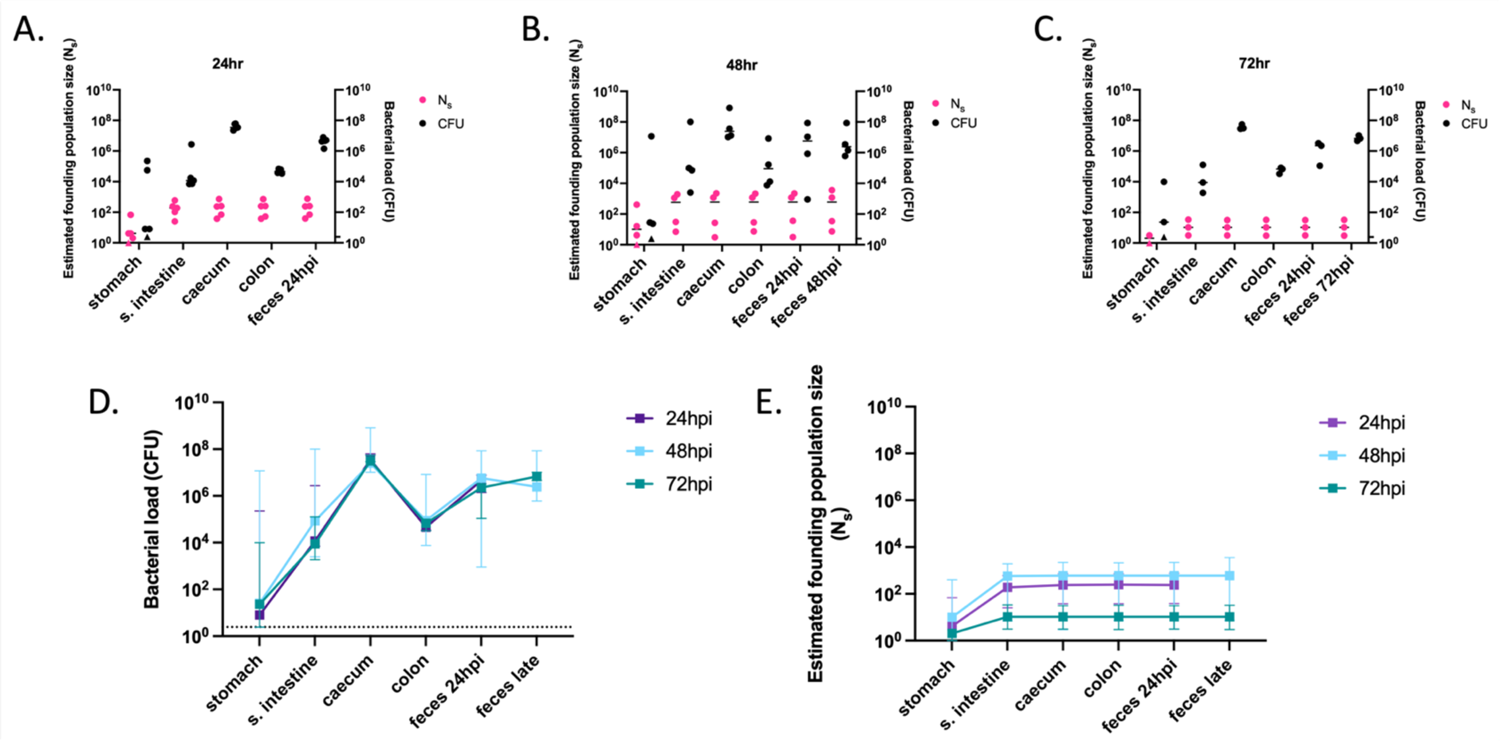
Founding populations and bacterial loads of *P. aeruginosa* in the GI tract. A total of 10^6.1^ CFU of PABL012_pool_ were delivered to single-caged mice by orogastric gavage. *P. aeruginosa* CFU in 250 µL of resuspensions of collected homogenized tissues (one-fourth of tissue homogenates of stomach, small intestine [“s. intestine”], caecum, colon, and feces) were enumerated by plating, and founding population sizes (N_s_) were estimated using the STAMPR approach. (A-C) Bacterial loads (CFU, black circles) and estimated founding population sizes (N_s_, pink circles) were quantified at (A) 24 (n = 5), (B) 48 (n = 4) and (C) 72 hpi (n = 3, except for N_s_ in stomach, which was n = 2 due to a sequencing issue). Each circle represents an organ from one mouse. Solid horizontal lines indicate medians. Minor ticks on the right Y axis represent the limits of detection for the CFU. Triangles represent samples with no recovered CFU. (D) *P. aeruginosa* burdens and (E) estimated founding population sizes in different tissues of the GI tract at 24 (purple), 48 (blue) and 72 (green) hpi. For comparison, fecal samples were collected at 24 hpi (“feces 24 hpi”) regardless of the ending timepoint. An additional terminal fecal sample was available for animals harvested at 48 or 72 hpi (“feces late”). Squares represent medians, and error bars represent 95% confidence intervals. The dotted line indicates the limit of detection for CFU. N_s_ values are not significantly different over time (t-test).

In all organs at all time points, N_s_ values were low, indicating that a tight bottleneck was encountered by *P. aeruginosa* following inoculation (Fig. 5A-C, E). N_s_ values were the lowest in the stomach, with median values below 10 for all three time points. Therefore, nearly all the bacteria initially inoculated into the stomach were either killed or expelled to the small intestine within the first 24 hours. The higher N_s_ values observed in the distal GI tract suggest that certain clones passed through the stomach but successfully established themselves further along the GI tract. By looking at the inoculum passage through the GI tract at early timepoints (1h and 6h post gavage), we confirmed that *P. aeruginosa* bacteria were mostly killed in the proximal GI tract rather than rapidly passaged to the distal GI tract and expelled in the feces (Supplemental Fig. 3). The nearly identical founding population sizes at each time point in the distal GI tract indicate that these segments are quite permissive for *P. aeruginosa* carriage (Fig. 5E). Taken together, these data suggest that nearly all *P. aeruginosa* bacteria are rapidly (in less than 6 hours) eliminated from the stomach and that a small number of bacteria pass through to the small intestine and downstream segments of the GI tract.

### The small intestine and caecum support high replication of *P. aeruginosa*

The large CFU counts and corresponding small founding populations in different organs highlighted the ability of *P. aeruginosa* to replicate in the GI tract (Fig. 5). For each segment of the GI tract, we defined net replication (which includes the combined effects of replication, death, and migration) as the ratio between CFU and N_s_. The greatest expansion of *P. aeruginosa* from a small founding population occurred in the caecum with CFU/N_s_ ratio greater than 10^5^, but substantial expansion was also observed in the stomach, small intestine and colon (Supplemental Fig. 4). Theoretically, high CFU/N_s_ could occur solely by local bacterial multiplication, by migration of bacteria *en masse* from an adjacent portion of the GI tract, or a combination of these two processes. Local multiplication would yield compartmentalized regions of the intestine, where different clones are spatially segregated along the length of the GI tract. In contrast, movement of bacterial populations along the GI tract would yield more similar barcode distributions between regions of the intestine. To distinguish between these possibilities, we quantified the genetic relatedness of *P. aeruginosa* populations in each segment of the GI tract. Genetic relatedness is determined by comparing the genetic distance (GD) of barcode distributions between two populations (45). GD varies from 0 to 0.9, with low values indicating highly similar barcode distributions between two samples and high values indicating different barcode distributions between samples. For all time points, the caecum, the colon, and the feces contained, on average, highly similar barcode populations of *P. aeruginosa* (GD ≤ 0.06) (Fig. 6 A-D, dark purple). Additionally, the GD between the small intestine and the caecum, colon and feces decreased over time (Fig. 6D, teal). These findings, along with the high CFU/N_s_ values observed in the distal GI tract (Supplemental Fig. 4), are consistent with a model in which *P. aeruginosa,* despite not being viewed as an enteric bacterium, multiplies rapidly and to high numbers in the caecum or the small intestine. These large populations of bacteria then move to the colon and are subsequently expelled in the feces. They also support the conclusion that *P. aeruginosa* bacteria recovered from fecal samples are most representative of those carried in the distal GI tract and confirm the utility of using fecal sampling to study *P. aeruginosa* carriage.

**Figure 6:**
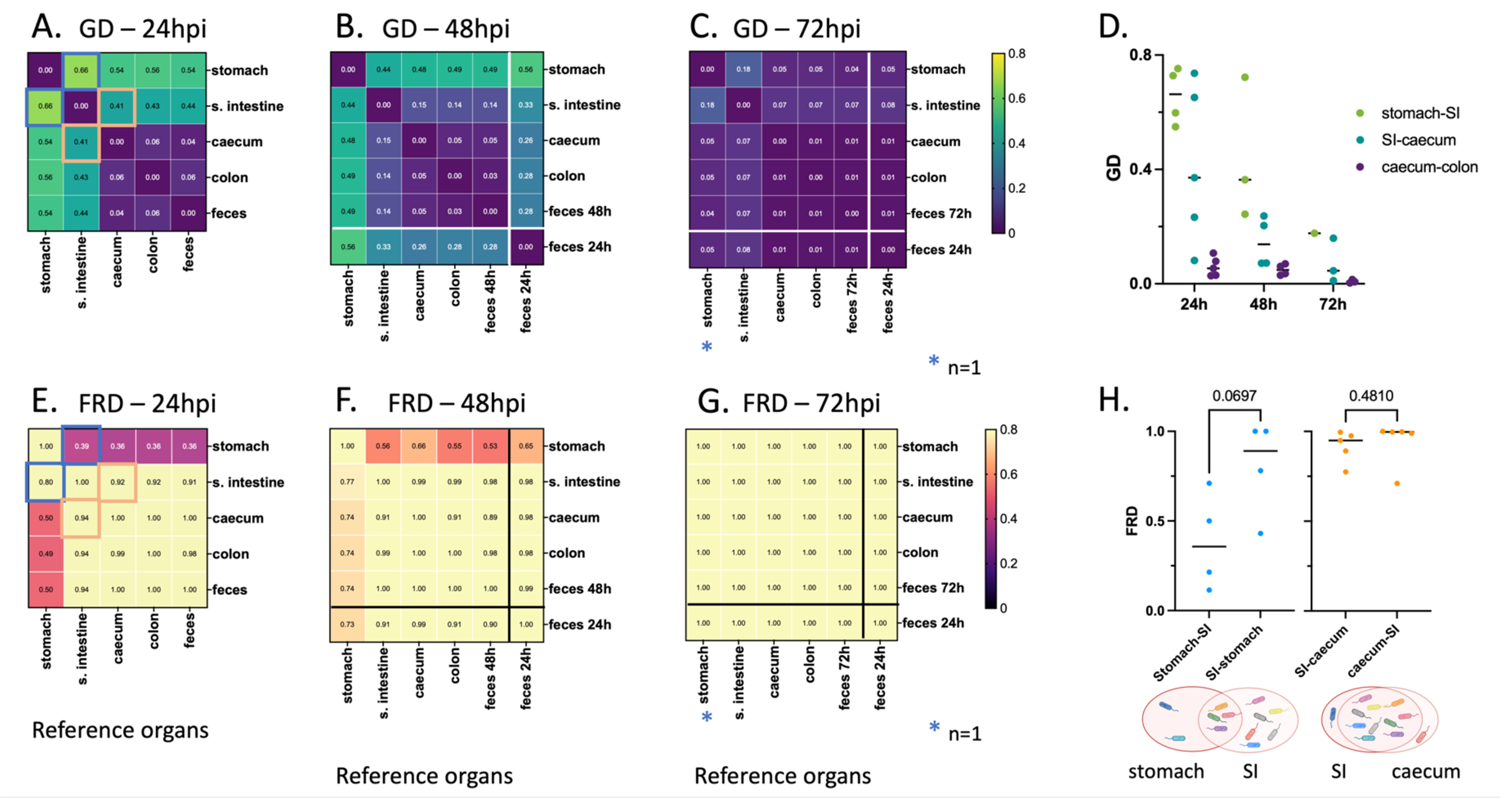
Average intra-mouse genetic relatedness of *P. aeruginosa* populations in the GI tract. (A-C) Heatmaps representing the average intra-mouse genetic distances (GDs) of *P. aeruginosa* from organs of the mice described in figure 5, at (A) 24, (B) 48, and (C) 72 hpi. Lower values of GD (purple) indicate a higher frequency of barcode sharing between the samples, with 0 reflecting identical populations. (D) GD values over time between: stomach and small intestine (“SI”) (green), small intestine and caecum (teal), and caecum and colon (purple). Each symbol represents one mouse. Lines indicate medians. (E-G) Heatmaps representing the average intra-mouse Fractional Resilient Genetic Distances (FRD) of *P. aeruginosa* from organs of the mice described in figure 5, at (E) 24, (F) 48 and (G) 72 hpi. The FRD is calculated using the following formula: 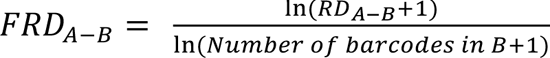 where RD_A-B_ is the number of shared barcodes that contribute to genetic similarity between samples A and B. The column names in the FRD heatmaps correspond to the organ of reference (B in the above formula). High FRD values (yellow) indicate that most bacterial barcodes are shared between samples. Thick lines in panels B-C, F-G separate the samples collected at the time of dissection (top/left) from samples of feces collected from the same animals at an earlier time point (“feces 24h”) (bottom/right). Samples outlined by blue and orange squares in panels A and E indicate pairs that are detailed in panel H. (H) FRD values for bacteria from the stomach/small intestine (SI) (blue) and small intestine/caecum (orange) pairs at 24 hpi. Each symbol represents one mouse. Lines indicate medians. p-values are indicated (two-tailed paired t test). The Venn diagrams under the graph are visual representations of the averaged proportion of barcodes shared between two adjacent organs (circles). Diagrams created using Biorender.com. As observed in figure 5, no bacteria could be detected in the stomach of some mice, leading to variation in the number of samples used for this analysis: A, E, H: n = 5 (except for the stomach; n = 4), B, F: n = 4 (except for the stomach; n = 3), C, G: n = 3 (except for the stomach; n = 1), D: see panels A-C.

Genetic similarity between two sites can be achieved through different population patterns. For example, two GI segments can be highly similar owing to a single dominant clone shared between both sites or due to underlying sharing of hundreds of clones, each with low abundance. To define the number of clones that contribute to genetic similarity, we calculated a metric known as “resilient” genetic distance (RD), which quantifies the number of shared clones that contribute to genetic similarity between two samples (0.8 threshold, see Methods) (43). Genetically similar samples with many shared clones have high RD values, whereas genetically similar samples which share only a few clones have low RD values.

The interpretation of whether RD is “low” or “high” is relative to the number of barcodes in a sample. To normalize RD values, the natural logarithm (ln) of RD is divided by the ln of the number of distinct barcodes, creating a fractional RD (FRD). The FRD represents the number of shared barcodes in a pair of samples (samples A and B) relative to the number of distinct barcodes in a reference sample (sample B) (43). For example, FRD_A-B_ is calculated as 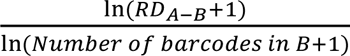, measuring the ratio of shared barcodes between samples A and B relative to the number of distinct barcodes in sample B. Therefore, an FRD_A-B_ of ≈ 1 indicates that nearly all barcodes shared between sample A and B are found in sample B, while low FRD_A-B_ indicates that shared clones represent a low fraction of barcodes in sample B. When contextualized with GD, FRD provides a normalized metric to interpret the number of shared clones that contribute to similarity and to suggest the possible directionality of clone transfer.

We next compared GD and FRD to decipher how clonal sharing of *P. aeruginosa* along the intestine changes over time. At 24 hpi, the *P. aeruginosa* populations in the stomach and the small intestine had only moderate similarity to those of the caecum, large intestine, and feces (average GDs = 0.54 and 0.41, respectively) (Fig. 6A). Using FRD values, we identified distinct patterns driving the genetic distances between the segments of the GI tract. At 24 hpi, the stomach and the small intestine had moderate genetic distance (average GD = 0.66) (Fig. 6A, D), indicating that the populations between these environments were relatively different. The median FRD_stomach-small intestine_ (0.36) was lower than FRD_small intestine-stomach_ (0.89) (Fig. 6H), illustrating that the shared barcodes constitute a smaller proportion of the total barcodes in the small intestine than in the stomach; the small intestine possesses a greater number of unique clones. The greater number of unique clones in the small intestine suggests either (1) reflux of a subpopulation of bacteria from the small intestine to the stomach or (2) initial seeding of the stomach and small intestine with more similar populations followed by rapid elimination of a portion of the population in the stomach. The decreasing GD between the stomach and the small intestine over time supports the idea of bacterial reflux from the small intestine. Very little similarity was present between *P. aeruginosa* populations in the feces and the stomach, indicating that coprophagia did not significantly contribute to reseeding of the stomach (Supplemental Fig. 5). Both the average FRD_small intestine-caecum_ and FRD_caecum-small intestine_ were greater than 0.9, indicating that relatedness between the small intestine and caecum is driven by a large portion of shared barcodes (Fig. 6E, H). The differential expansion of a small number of clones likely accounted for the genetic distance (average GD = 0.41) between these two sites (Fig. 6A, Supplemental Fig. 5). The high FRD and low GD values between the small intestine and the caecum suggest that *P. aeruginosa* clones efficiently trafficked between these two compartments, either through natural peristalsis or retrograde movement, before continuing expansion at both sites. On the other hand, the extremely low GD values between the caecum, colon, and feces (Fig. 6A-C) with corresponding FRD values ≥ 0.89 (Fig. 6E-G) suggest that bacterial populations move freely between these sites. These observations, together with the fact that CFU are consistently higher than N_s_ values, indicate that vancomycin pretreatment robustly enables *P. aeruginosa* replication and movement along the GI tract, rather than simply facilitating the transient transfer of an initial inoculum through the GI tract.

Overall, our findings suggest that (i) the vast majority of orogastrically administered *P. aeruginosa* bacteria are rapidly (within 6 hours) killed in the stomach, (ii) less than 0.01% of *P. aeruginosa* from the inoculum persists in the intestine over the first 72 hours, (iii) robust *P. aeruginosa* replication occurs in the small intestine and the caecum, and (iv) bacterial populations subsequently migrate along the distal GI tract and are expelled in the feces.

## DISCUSSION

In this study, we established a murine model of *P. aeruginosa* GI carriage that mimics patients receiving antibiotics in the hospital setting. This model is clinically relevant, as it utilizes vancomycin, and has been validated with both sexes and multiple clinical isolates. The absence of GI tract inflammation confirmed that this model represents carriage, not infection. Nevertheless, occasional low-level escape of *P. aeruginosa* to other organs suggests a possible route by which GI carriage may lead to subsequent infection at remote sites (7, 12–14). Long-term GI carriage was established after a single dose of *P. aeruginosa*, with 70% of mice still carried the bacterium after 60 days. Using barcoded bacteria, we found that most of the bacterial inoculum was eliminated within the first 6 hours, primarily in the stomach. However, once *P. aeruginosa* reached the small intestine and the caecum, bacteria replicated robustly, leading to significant fecal excretion as winnowed bacterial populations migrated unimpeded through the caecum and colon. Confirming previously reported results (28), we found that untreated mice did not support GI carriage of *P. aeruginosa.* In contrast, seven days of vancomycin delivered through IP injection promoted high-level and prolonged GI carriage of *P. aeruginosa*.

Several studies have investigated the impact of orally administered vancomycin on the gut microbiota and have reported a decrease in *Bacteroidetes* and a subsequent increase in *Proteobacteria* and *Fusobacteria* phyla (46–50). *Firmicutes* levels were also altered by oral vancomycin treatments, with the directionality of the impact varying across bacterial species. We speculate that IP delivered vancomycin achieves relatively high concentrations in the lumen of the GI tract, that it has a similar effect on the mouse microbiome, and that depletion of some microbiome constituents from the GI tract creates a niche for the establishment of *P. aeruginosa*. While we did not observe histopathological changes following vancomycin treatment, it is also possible that this antibiotic facilitates *P. aeruginosa* carriage through a direct effect on the host, such as immunomodulation, independent from a modification of the microbiome (49). As a first step in understanding the mechanism by which vancomycin facilitates *P. aeruginosa* carriage, we are currently characterizing the changes in the microbiome over time following administration of vancomycin in this model.

All six clinical isolates of *P. aeruginosa* tested in our study established carriage in the GI tract to a similar extent. These isolates are genetically diverse and include both *exoU^+^* and *exoS^+^* strains, as well as high-risk and non-high-risk clones (38). Robust colonization by different isolates suggests that the ability for *P. aeruginosa* to establish GI carriage depends on a set of features encoded by the core genome of the bacterium. Genes encoding metabolic factors, transcriptional regulators, adhesins, secretion systems, membrane homeostasis proteins and bile resistance factors have been identified as important for the GI carriage of other bacterial species (51–55). We suspect that *P. aeruginosa* carriage requires a similar set of features. The reliance on additional strain-specific strategies is nonetheless not excluded. Treatment with vancomycin for seven days likely caused a severe dysbiosis, masking the need for strain-specific factors that contribute to carriage in the presence of a less perturbed microbiome. Additionally, we observed that GI carriage can last for several weeks following a single exposure to the bacterium and cessation of vancomycin. Several studies have reported that the long-term carriage of *P. aeruginosa* in the lungs of individuals with cystic fibrosis or chronic obstructive pulmonary disease is accompanied by genetic adaptations of the bacterium (56–58). It is possible that the set of genes or alleles contributing to carriage of *P. aeruginosa* evolves as bacteria transition from the early stages of carriage to long-term colonization.

In patients, the GI carriage of *P. aeruginosa* is a predictor for the subsequent development of *P. aeruginosa* infections at various sites (7, 12, 13). Shortly after bacterial inoculation, we show that low-level dissemination from the gut to the gallbladder, spleen, liver, or lungs occasionally occurs, perhaps explaining these clinical observations. Gut dysbiosis can lead to the development of a leaky intestinal barrier, where pathogen molecules translocate from the gut into the bloodstream (59). The present study utilized healthy mice. However, dissemination to other tissues may be accentuated in immunocompromised animals, leading to more severe infections, as shown by Koh *et al.* (18). Interestingly, 12 days after cessation of vancomycin administration (14 days after orogastric gavage with *P. aeruginosa*), *P. aeruginosa* CFU in the feces remained high but dissemination from the gut was no longer observed. Calderon-Gonzalez *et al.* showed that *Klebsiella pneumoniae* dissemination from the GI tract was promoted by antibiotic treatments (51). It is therefore possible that vancomycin may support not only GI carriage but also dissemination of *P. aeruginosa,* and that dissemination diminishes over time as vancomycin is cleared.

STAMPR has been used by several groups to measure the dynamics of bacterial spread in colonization and systemic dissemination (44, 60, 61). We used STAMPR to show that most *P. aeruginosa* bacteria are eradicated prior to carriage. Two recent studies indicate that stomach acidity significantly restricts the GI carriage of *Citrobacter rodentium* and *K. pneumoniae* (51, 62). While stomach acidity constricts *C. rodentium* numbers by 10- to 100-fold (62), our results show an even more severe constriction for *P. aeruginosa* in the stomach. The stomach pH of healthy mice fluctuates between 3-4, while the rest of the GI tract tends to have a more neutral pH of 6-8 (63). *P. aeruginosa* can grow at a wide range of pH, but its optimal growth pH ranges between 6 and 8 (64). Thus, acidity may be the mechanism underlying the loss of *P. aeruginosa* barcode diversity in the stomach. This possibility could be tested by pharmacologically neutralizing stomach acid and subsequently measuring the size of downstream founding populations of *P. aeruginosa.* Interestingly, the rest of the GI tract was quite permissive to carriage. In the absence of vancomycin treatment, no carriage could be established, indicating the presence of a second barrier to *P. aeruginosa,* perhaps downstream of the stomach in untreated mice. Campbell *et al.* recently monitored the dynamics of *C. rodentium* enteric carriage and identified the microbiota as the major factor limiting colonization (62). This supports the idea that vancomycin facilitates the carriage of *P. aeruginosa* by eliminating a microbiome-mediated barrier, a hypothesis that could be tested with germ-free mice. In this sense, *P. aeruginosa* is both an “opportunistic pathogen” and an “opportunistic colonizer.”

Together, our findings suggest the following model of *P. aeruginosa* carriage: within a few hours (less than 6 hours), a drastic constriction (less than 0.01% survival on average) of the bacterial inoculum occurs in the stomach (Fig. 7). A very small proportion of the *P. aeruginosa* inoculum passes through the stomach to reach the small intestine and the caecum. The small number of remaining founders rapidly replicate in both the small intestine and the caecum, and the resulting bacterial populations migrate from the caecum to the colon and are expelled in the feces. In addition to the expected trafficking route from the stomach to the small intestine, a small portion of *P. aeruginosa* may also reflux from the small intestine back to the stomach. Using STAMPR, we have identified the stomach as the main barrier to *P. aeruginosa* carriage in this animal model and the small intestine and the caecum as the main sites of bacterial expansion.

**Figure 7:**
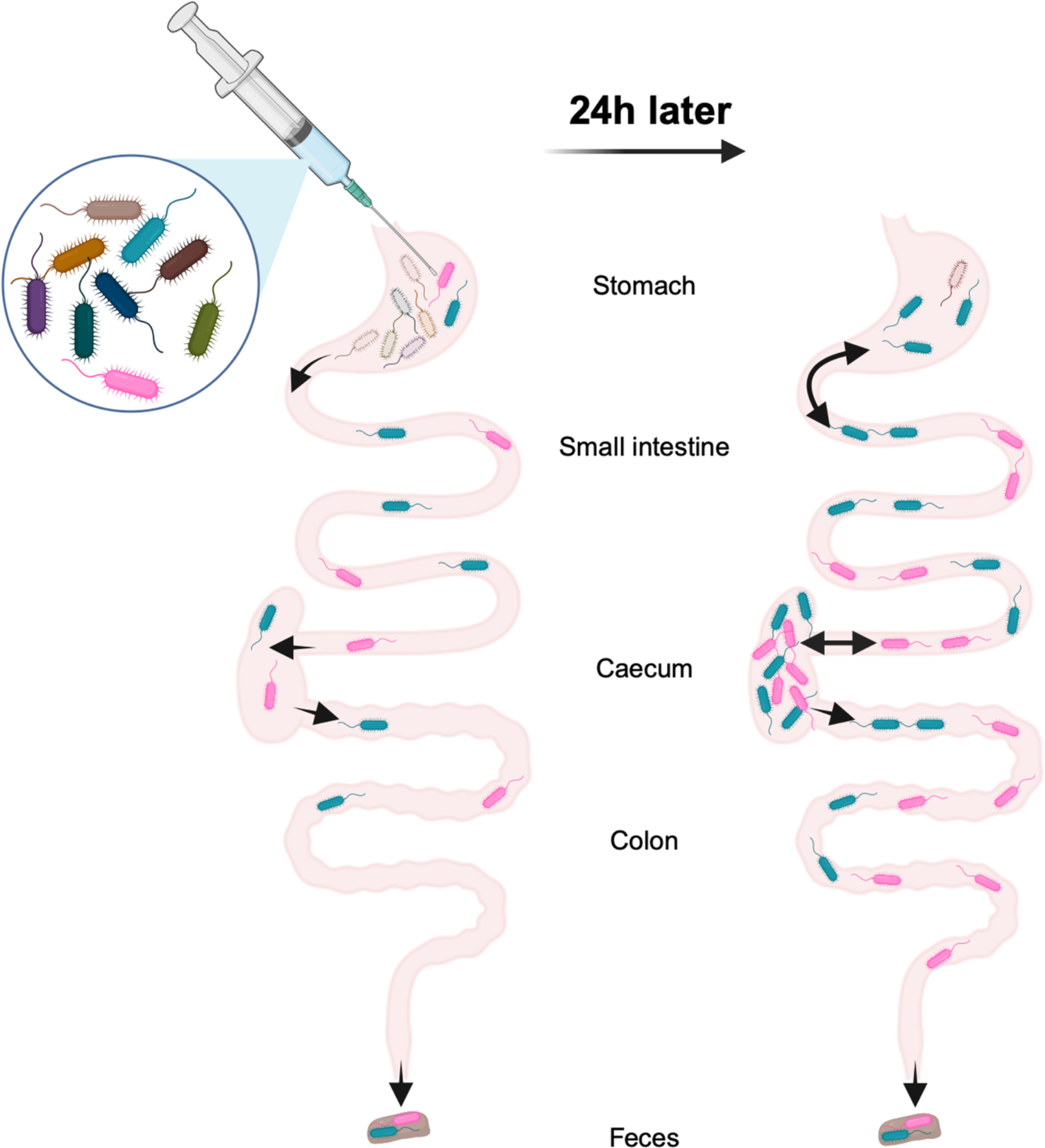
Model of the population dynamics of *P. aeruginosa* following orogastric gavage. Left: soon after the orogastric delivery of *P. aeruginosa*, most bacteria are eliminated from the stomach, severely constricting the size of the remaining population (less than 0.01% survival). Part of the population passes through the stomach to reach other compartments of the GI tract: small intestine, caecum, colon, and feces. *P. aeruginosa* does not encounter additional barriers downstream from the stomach. Right: over the first 24 hours, population expansion and/or reflux from the small intestine occurs in the stomach. The small intestine and the caecum support massive expansion of the remaining *P. aeruginosa* clones, and bacteria freely migrate from the caecum to the colon and feces. This figure was created using Biorender.com.

Overall, this model advances the understanding of *P. aeruginosa* dynamics during GI carriage and may be useful in studying adaptation of *P. aeruginosa* during prolonged colonization, persistence following administration of antibiotics other than vancomycin, and identification of *P. aeruginosa* and host factors that facilitate carriage. In addition, our findings have clinical implications, such as the potential importance of more judicious use of acid suppressing drugs and vancomycin in preventing *P. aeruginosa* GI carriage.

## MATERIALS AND METHODS

### Bacterial strains and culture conditions

PABL004, PABL006, PABL012, PABL048, PABL049 and PABL054 are archived *P. aeruginosa* clinical isolates cultured between 1999 and 2003 from the bloodstream of patients at Northwestern Memorial Hospital in Chicago (65). Relevant characteristics of the strains are listed in Supplementary Table 1. Unless otherwise stated, bacteria were streaked from frozen stocks onto either Lysogeny broth (LB) or Vogel-Bonner minimal (VBM) (66) agar plates and subsequently grown at 37°C in LB medium with shaking. When antibiotic selection for *P. aeruginosa* was necessary, supplementation with irgasan (irg) at 5 µg/mL was used.

### Murine model of gastrointestinal carriage

Six- to eight-week-old C57BL/6 mice (Jackson Laboratory) received either 200 µL (female) or 250 µL (male) of vancomycin (370 mg/kg, Hospira, Lake Forest, IL) daily IP for seven days. The vancomycin dosage was allometrically scaled based on a total human daily dose of 30 mg/kg (35). On the fifth day of antibiotic treatment, mice were gavaged (20 G x 30 mm straight animal feeding needle, Pet Surgical, Phoenix, AZ) with 50 µL of *P. aeruginosa* prepared as follows: after overnight culture in LB, bacteria were diluted, regrown to exponential phase in LB and resuspended to the desired dose in PBS. When specified, mock IP injections or mock orogastric gavage was performed using PBS (same volumes as the treatment groups). Starting the day after the orogastric gavage, cages were changed daily to limit the impact of coprophagy.

To determine bacterial GI carriage, mice were individually placed in boxes to induce defecation, and feces were collected, weighed, homogenized in 1 mL of PBS using a bead blaster (Benchmark Scientific, Sayreville, NJ), and centrifuged for 30 sec at 1,100 x *g*. The supernatant was serially diluted and plated on either LB agar supplemented with irgasan or VBM agar for CFU enumeration.

Mice were housed in the containment ward of the Center for Comparative Medicine at Northwestern University. All experiments were approved by the Northwestern University Institutional Animal Care and Use Committee in compliance with all relevant ethical regulations for animal testing and research. Experiments used female mice unless otherwise stated.

### Murine GI carriage of PABL012_pool_

Six- to eight-week-old female mice received IP injection of vancomycin (200 µL, 370 mg/kg) for 5 to 7 days, with each experimental group receiving their last injection 24 h prior to dissection. An aliquot of 50 µL of the PABL012_pool_ library was grown overnight in 5 mL of LB (37°C, 250 rpm) and subcultured (1:40) in 30 mL of LB for 3 h. The bacterial inoculum was prepared as described above, and orogastric gavage was performed on the 5^th^ day of vancomycin treatment, using 10^6.1^ CFU of PABL012_pool_.

Following bacterial inoculation, mice were housed individually. At 24, 48, and 72 hours post-gavage, mice were euthanized, and the stomach, small intestine, caecum, colon and feces were collected. The stomach, small intestine and caecum were processed along with their luminal contents. The colon was emptied, and the colonic contents were added to excreted feces (when available) to constitute the “feces” samples. All samples were weighed, homogenized in 1 mL of PBS using a bead blaster, centrifuged for 30 sec at 1,100 x *g*, and the supernatant was serially diluted and plated on VBM agar for CFU enumeration. For the estimated founding population sizes (N_s_), 250 µL of the organ samples, as well as 250 µL of the inoculum (26 technical replicates) were spread on 150-mm-diameter VBM plates. Plates used for CFU and N_s_ determination were grown overnight at 37°C and the colonies were counted. CFU counts and N_s_ values in figure 5 represent those determined in 250 µL (1/4 of the homogenized tissue volume).

## ACKNOWLEDMENTS

Support for this work was provided by the National Institutes of Health awards RO1 AI118257, K24 AI04831, R21 AI129167 and R21 AI153953 (all to ARH) and U19 AI135964 (RGW and ARH). Histology services were provided by the Northwestern University Mouse Histology and Phenotyping Laboratory which is supported by NCI P30-CA060553 awarded to the Robert H Lurie Comprehensive Cancer Center.

## Supplemental Figure Legends

**Supplemental Figure 1:**
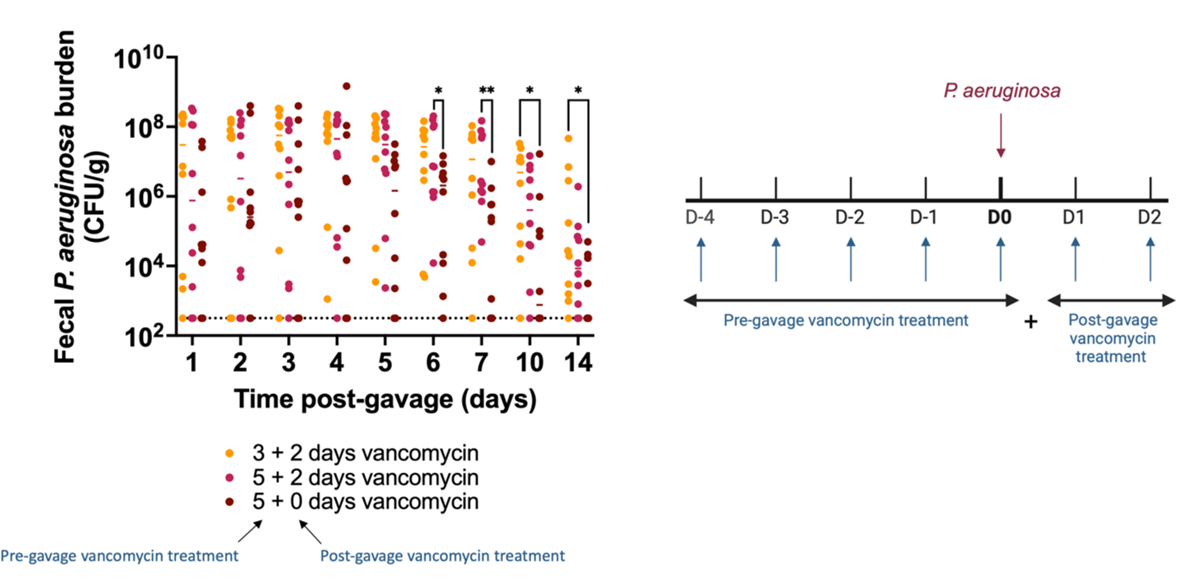
GI carriage of *P. aeruginosa* obtained with various regimens of vancomycin treatment. Mice received daily injections of vancomycin for various times before and after orogastric gavage (“x + y days” with x = the number of days of vancomycin injections prior to and on the day of orogastric gavage, and y = the number of days of vancomycin injections after the bacterial inoculation). Orogastric gavage was performed with 10^5.8+/-0.2^ CFU of strain PABL048. Each symbol represents one mouse. Solid horizontal lines indicate medians. The experiment was performed twice (combined results shown; n = 10)). The dotted line indicates the limit of detection. *p ≤ 0.05, **p ≤ 0.01 (t-tests). Significant differences were not detected for any of the time points between mice treated with 5 + 2 days and 3+ 2 days of vancomycin.

**Supplemental Figure 2:**
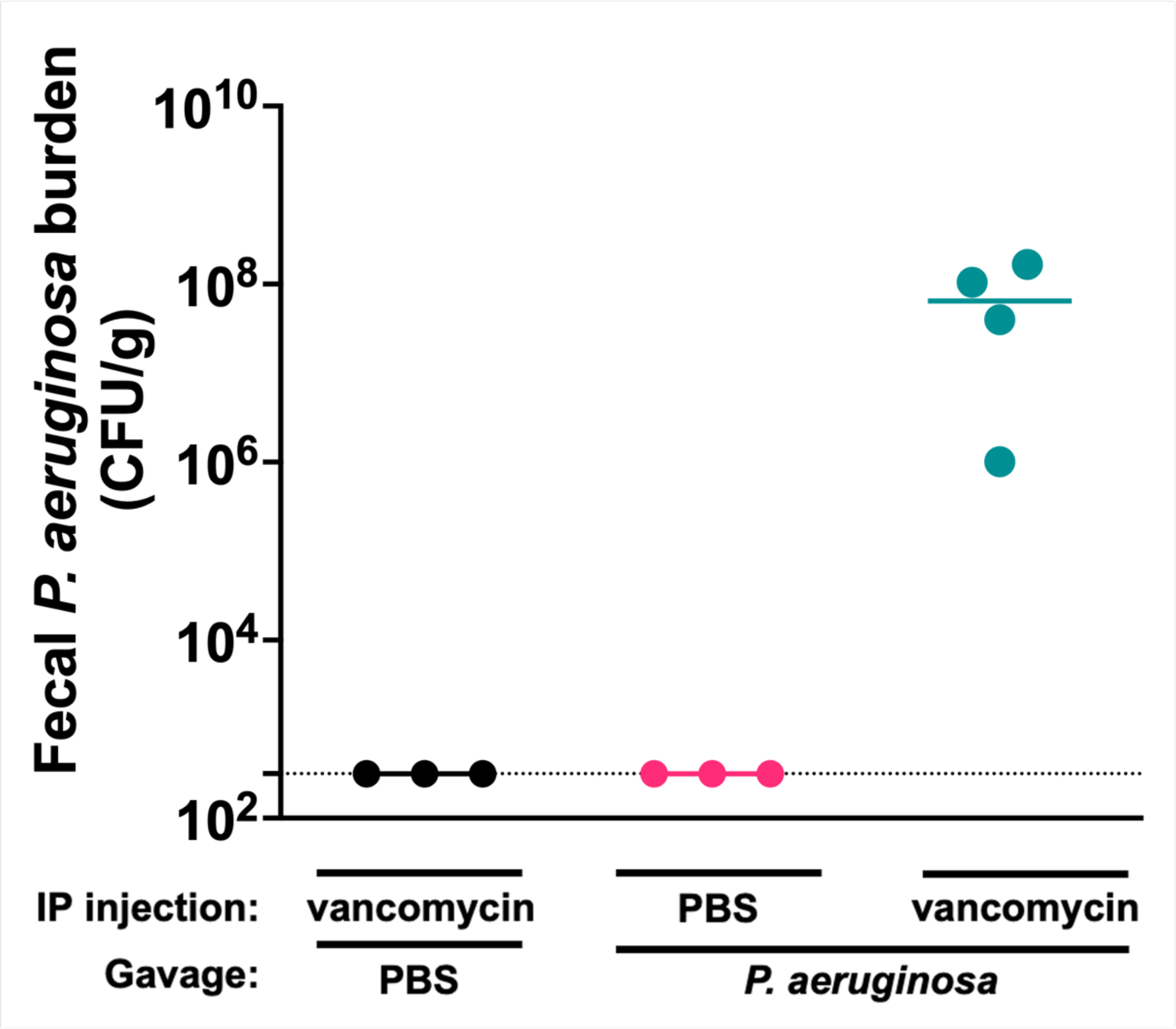
Fecal burden of strain PABL048 at day 3 post-inoculation. Mice were treated with either PBS (pink) or vancomycin (black and teal) for 7 days. On the fifth day of treatment, mice received either PBS (black) or 10^7.1^ CFU of PABL048 through orogastric gavage (pink and teal). The experiment was performed once (n=3-4 animals/group). Each symbol represents one mouse. Lines indicate medians. The dotted line indicates the limit of detection.

**Supplemental Figure 3:**
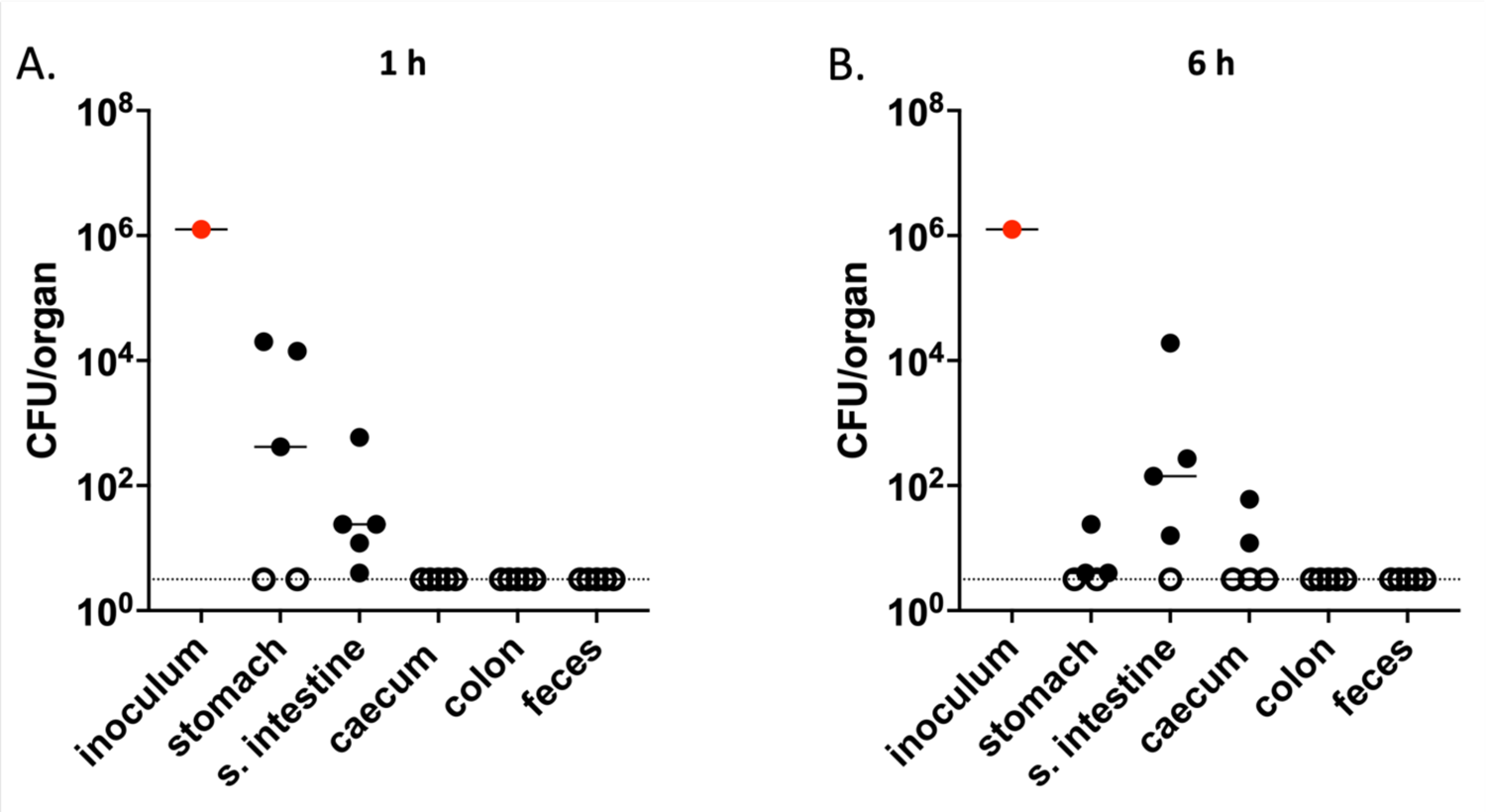
Recovery of *P. aeruginosa* from the GI tract at early times following inoculation. *P. aeruginosa* burden in GI tissues of mice gavaged with PABL012. Mice were sacrificed at (A) 1 h (n = 5) or (B) 6 h (n = 5) post-orogastric gavage with 10^6.1^ CFU of PABL012, and bacterial CFU in the organs were enumerated by plating. Experiment performed once. Red circles represent the inoculums. Each black circle represents one mouse. Solid horizontal lines indicate medians. The horizontal dotted line indicates the limit of detection. Open circles represent tissues with CFU bellow the limit of detection.

**Supplemental Figure 4:**
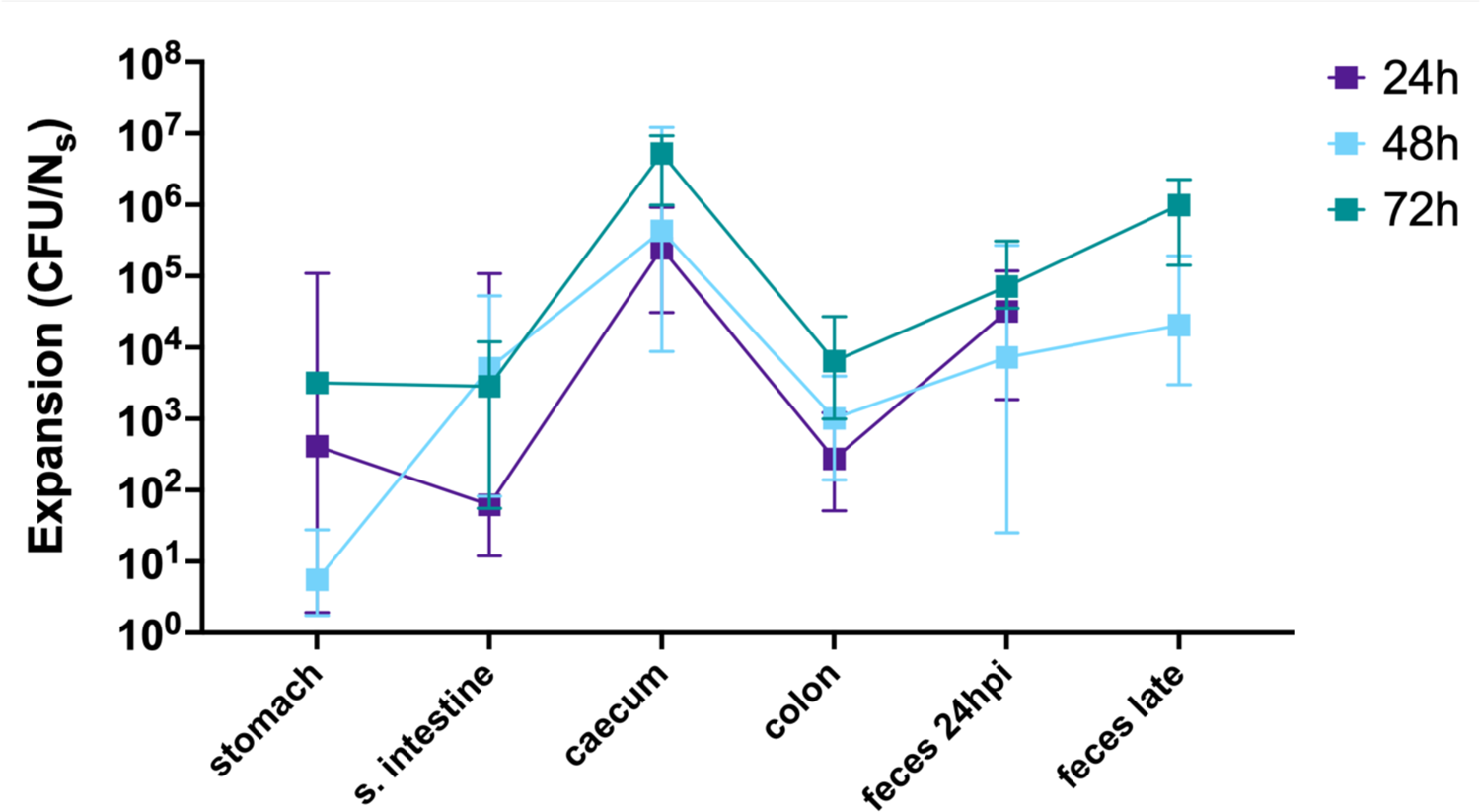
Ratio of bacterial recovery vs. founding population in GI sites. Tissues were harvested at 24 (purple, n = 5), 48 (blue, n = 4) or 72 hours (green, n = 3) after orogastric gavage with PABL012_pool_. Fecal samples were collected at 24 hpi (“feces 24 hpi”) regardless of the ending timepoint. Additional terminal fecal sample timepoints were available for animals that had organs harvested at 48 or 72 hpi (“feces late”). CFU/N_s_ ratios were calculated. Squares represent medians, and error bars represent the 95% confidence intervals.

**Supplemental Figure 5:**
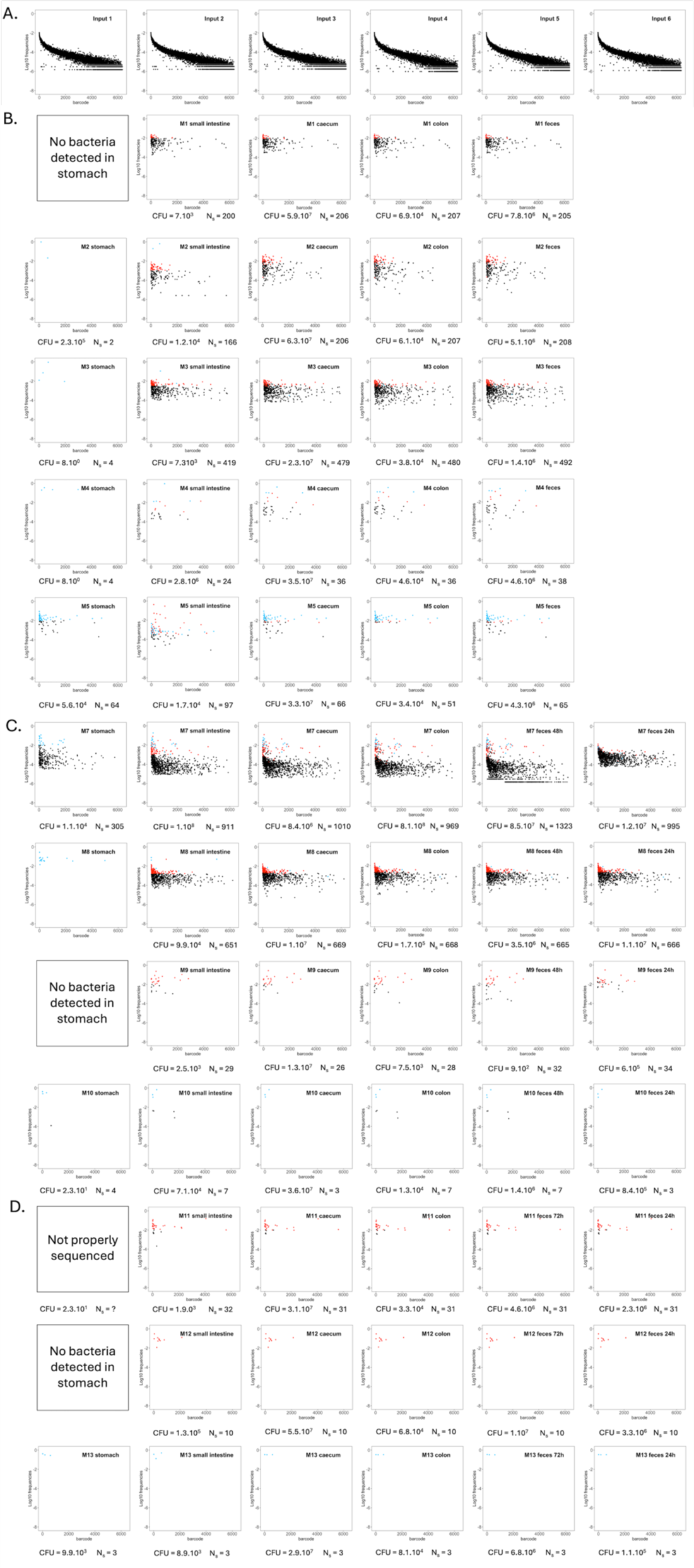
Barcode frequency distributions of *P. aeruginosa* bacteria recovered from mice following orogastric inoculation. The frequencies of unique barcodes in each bacterial population from different sites are shown. (A) Inoculum samples. Barcode frequency was analyzed in the 26 bacterial aliquots that were each used to inoculate a different mouse in the STAMP experiment. Six representative frequency distributions are shown. (B-D) Barcode frequency distributions after noise removal for the output samples from mice sacrificed at (B) 24, (C) 48 or (D) 72 hours post-orogastric gavage. Each dot represents the frequency at which one specific barcode was detected. For each mouse (“M#”), dots representing the most frequent clones identified in the stomach are colored blue in all organs, and dots representing the most frequent clones identified in the small intestine are colored red.

